## Supplementary material for "High resolution cryo-EM and crystallographic snapshots of the large actinobacterial 2-oxoglutarate dehydrogenase: an all-in-one fusion with unique properties": Suppl. Figures and References

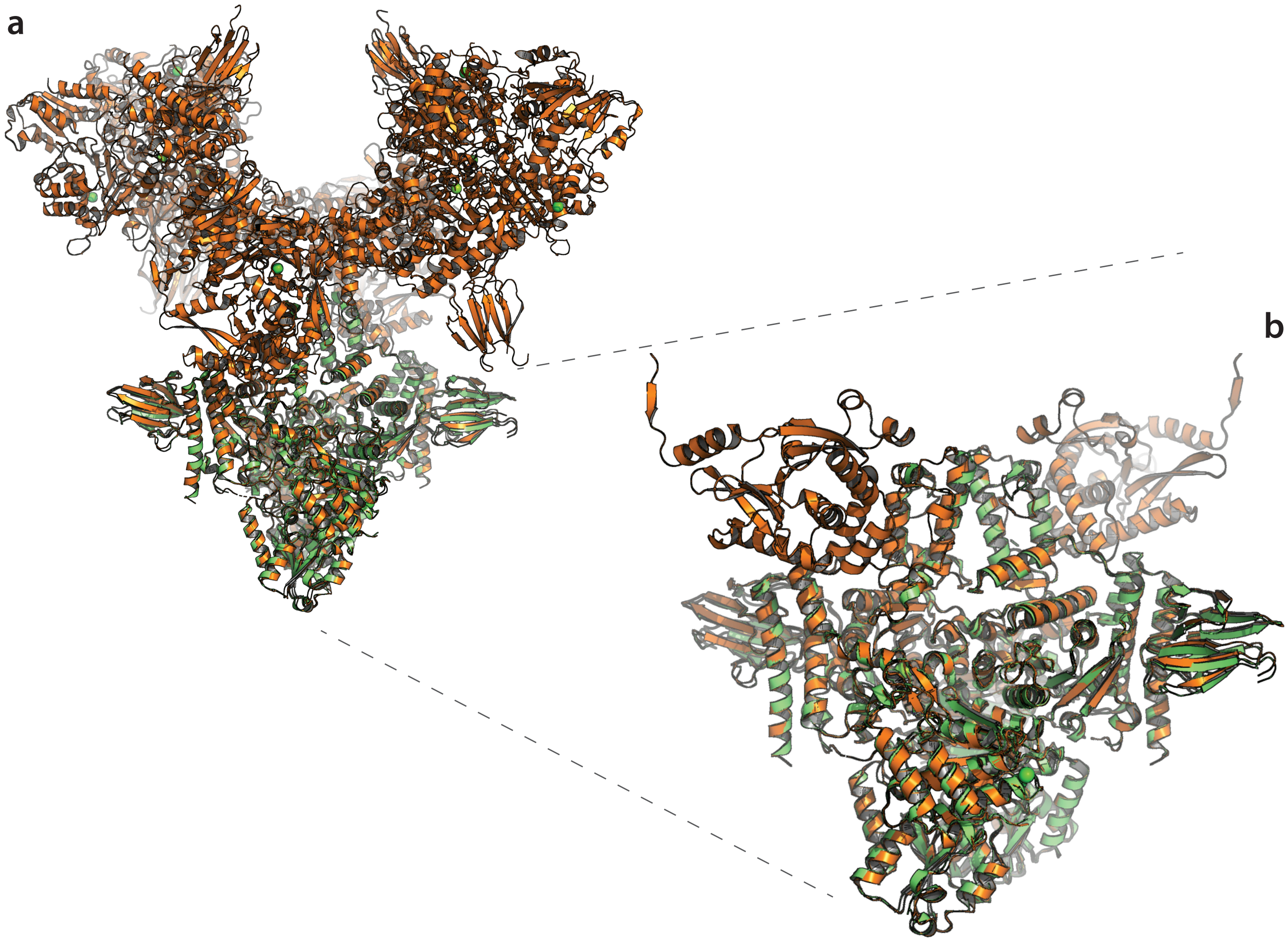


**Suppl. Figure 1**. ***Ms*KGD-GarA complexes.**
(**a**) Superimposition of homohexameric *Ms*KGD in complex with GarA (orange; this work) to the previously reported *Ms*KGD$_{\Delta 360}$-GarA complex (green; PDB 6I2Q [1]). For clarity, only one *Ms*KGD$_{\Delta 360}$ dimer, with bound GarA, is superimposed to the hexameric full-length *Ms*KGD$_{\Delta 360}$-GarA. (**b**) Zoomed view on a *Ms*KGD full-length dimer, isolated from its hexameric assembly, with superimposed *Ms*KGD$_{\Delta 360}$-GarA. Coordinates of *Ms*KGD$_{\Delta 360}$-GarA can be superimposed to full-length *Ms*KGD$_{\Delta 360}$-GarA with an RMSD of 0.6 Å (C$\alpha$ atoms).

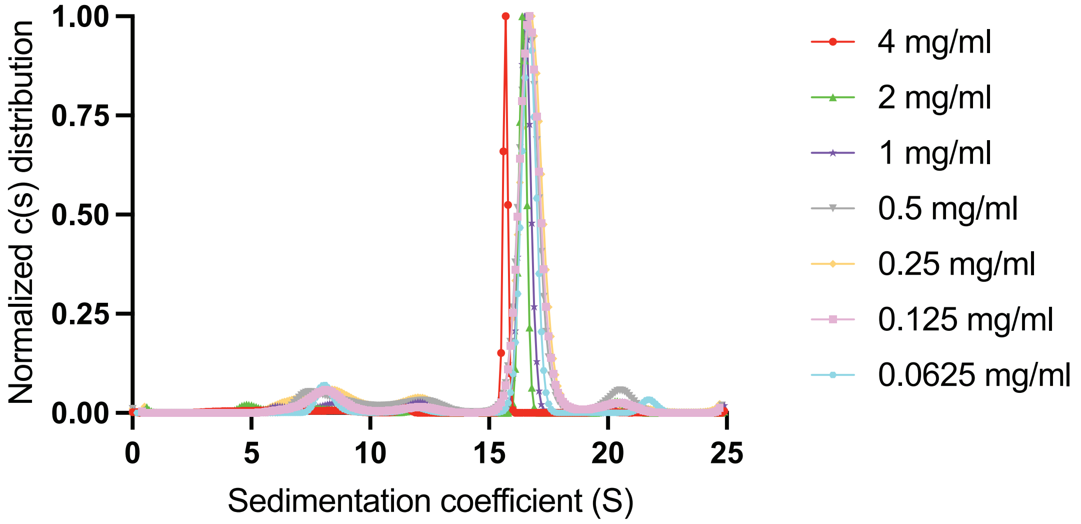


**Suppl. Figure 2. OdhA analytical ultracentrifugation profiles at decreasing protein concentrations.** Normalized sedimentation coefficient distributions c(s) are shown for each OdhA concentration tested.

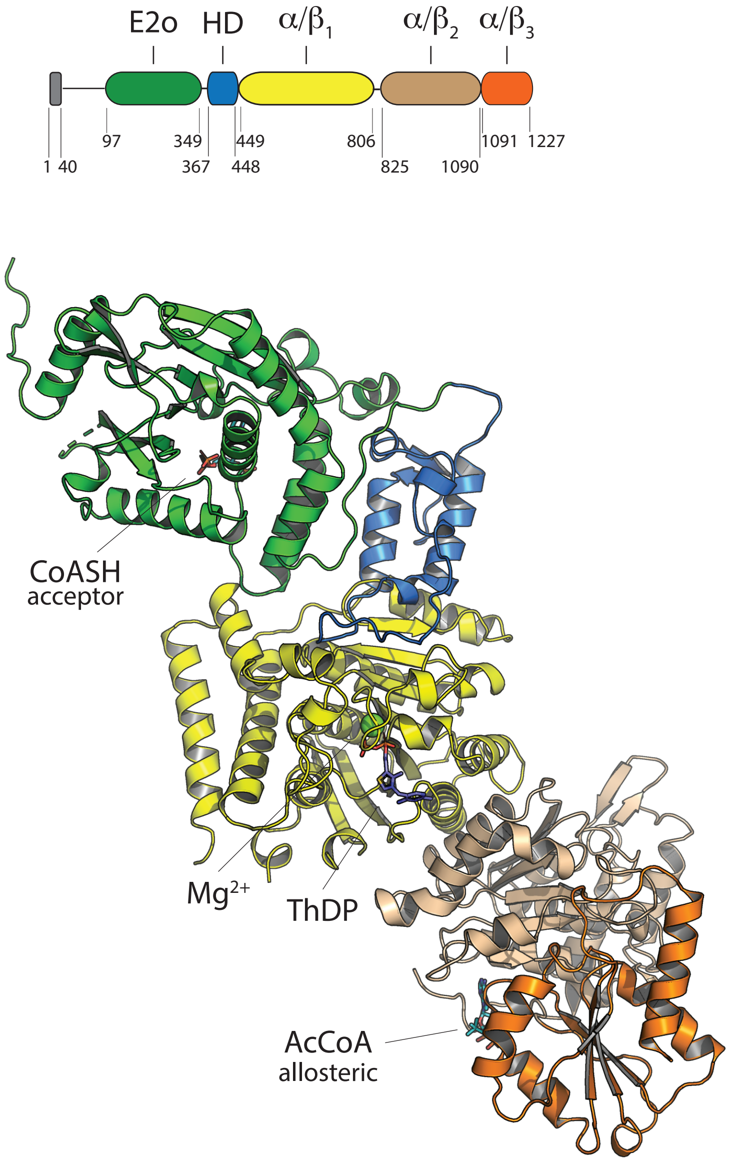


**Suppl. Figure 3. OdhA domain substructure.**
Cartoon representation of the OdhA$_{\Delta97}$ monomer, determined by X-ray crystallography. Colors according to the individual subdomains. HD: helical domain. The E1o catalytic domain is composed of the HD followed by three consecutive $\alpha/\beta$ domains.

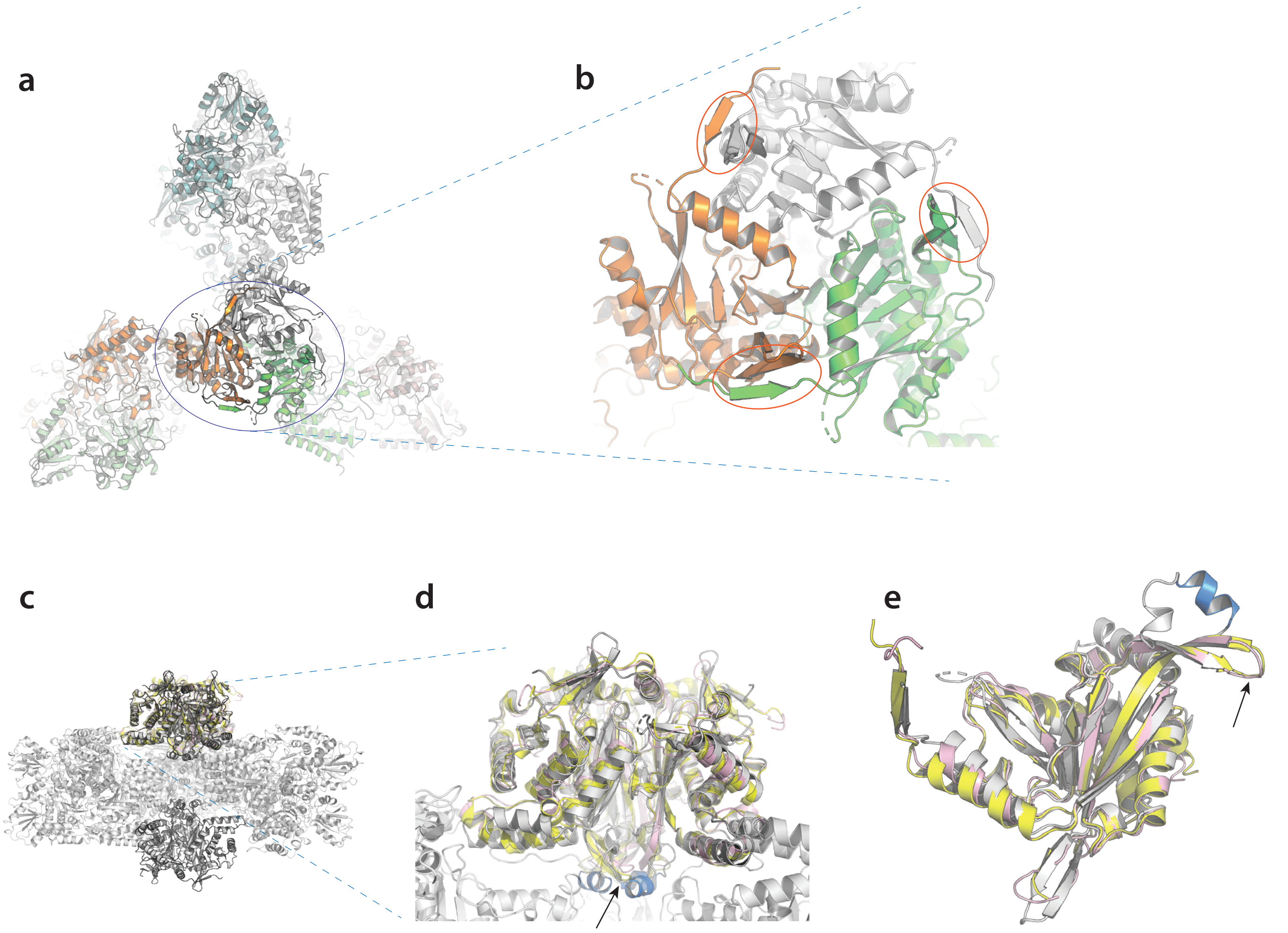


**Suppl. Figure 4. The OdhA N-terminal E2o domain.**

(**a**) Top view of the OdhA homohexamer in cartoon representation (one colour per chain). (**b**) Isolated view of the OdhA E2o domain (one colour per chain), with red ellipses indicating the mixed β-sheets composed by exchanged N-terminal strands. (**c**) Lateral OdhA view with E2o catalytic domains from *E. coli* (PDB 1SCZ; yellow) and *H. sapiens* (PDB 6H05 [2]; pink) superimposed to its N-terminal succinyltransferase domain. For clarity, only a single homotrimeric catalytic unit from both the *E. coli* and human E2o orthologs was superimposed (top); both proteins actually make 24-mer cubic assemblies with 432 symmetry. (**d, e**) Zoomed view on the superimposed E2o domains, with focus on a whole E2o trimer (d) or on superimposed single monomers (e). Colours as in (c). Despite the conserved, trimeric chloramphenicol acetyltransferase fold, a major difference is made, in OdhA, by the lack of the β-hairpin which, in canonical E2o enzymes, lies close to the 3-fold axis and is oriented towards the inside surface of the 24-mer cube (arrows). The corresponding region makes a short α-helix instead (Glu296-Leu302; blue), connected by a flexible linker, and which interacts with residues that belong to the E2o-E1o linker as well as to the N-terminal helical subdomain from the E1o domain (d).

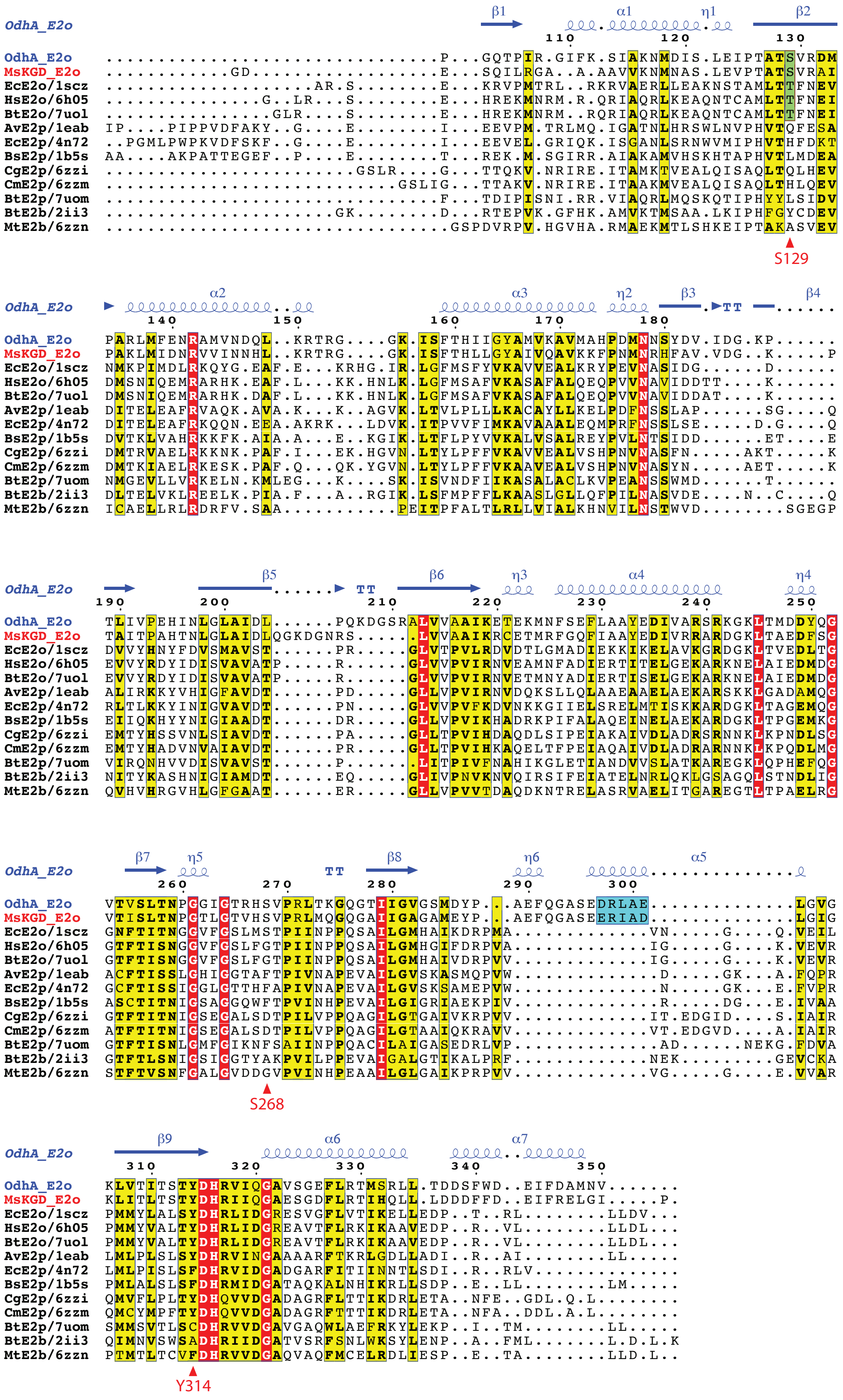


**Suppl. Figure 5. Multiple structural alignment of characterized acyltransferase (E2) domains.**
Sequence names start with two letters indicating the source organism (Av: *Azotobacter vinelandii*; Bs: *Bacillus stearothermophilus*; Bt: *Bos taurus*; Cg: *Corynebacterium glutamicum*; Cm: *Corynebacterium mustelae*; Ec: *Escherichia coli*; Hs: *Homo sapiens*; Ms: *Mycobacterium smegmatis*; Mt: *Mycobacterium tuberculosis*) and are followed by the respective four-character PDB entry codes. The sequence alignment is derived from a multiple structural alignment performed by PDBeFold [3], depicted with ESPript [4]. Red triangles on the bottom indicate residues involved, in OdhA, in H-bonding the succinyl moiety of the succinyl-CoA product. The blue boxing indicate residues making a short α-helix (here indicated as α5) part of a specific structural motif in both OdhA and *Ms*KGD (insertion corresponding to OdhA residues 289-301; see also Suppl. Figure 4).

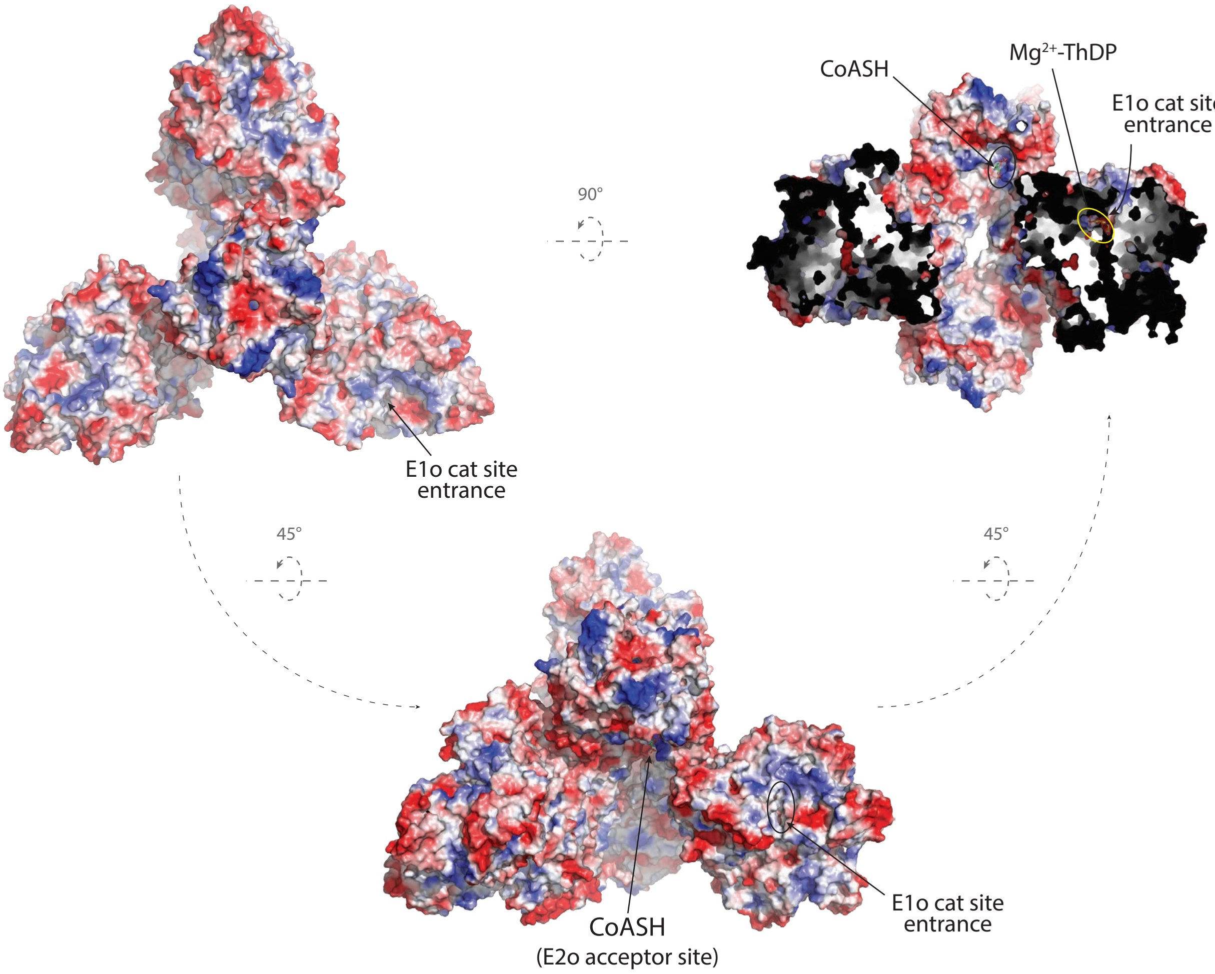


**Suppl. Figure 6. E1o and E2o active site localization and accessibility in the OdhA homohexamer.**
The OdhA$_{\Delta97}$ homohexamer is shown here with its surface coloured according to the eletrostatic surface potential (calculated and rendered with pymol [5]), depicted at different orientations following rotation of the hexamer plane. The positions of the E1o and E2o active site, and the respective ligands are indicated ($Mg^{2+}$-ThDP cofactor for the E1o site, CoASH acceptor for E2o).

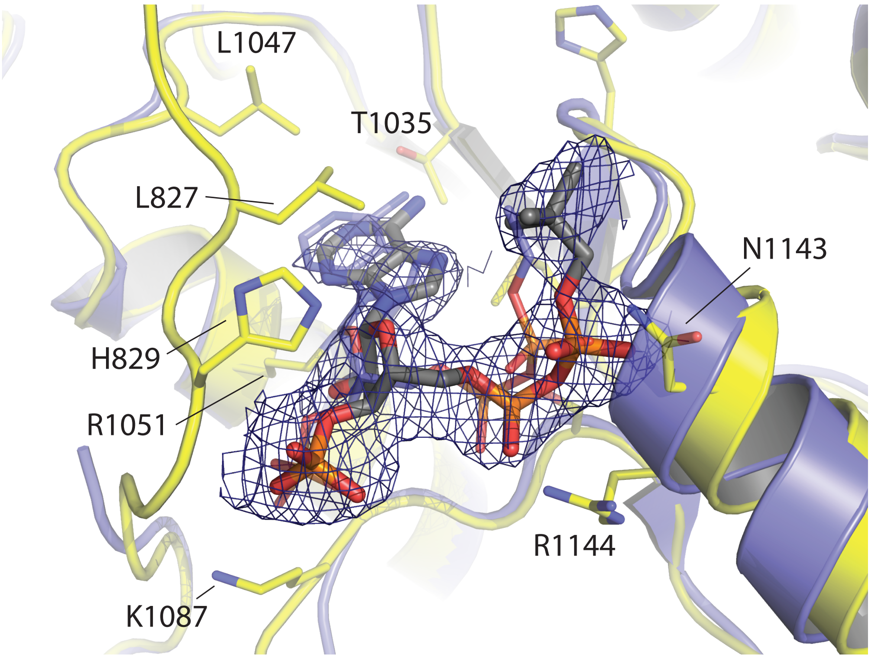


**Suppl. Figure 7. Acetyl-CoA allosteric pocket on OdhA$_{D97}$ (E1o domain).**
E1o allosteric site in the OdhA$_{D97}$ crystal structure, showing a bound ligand compatible with a CoA ester (presumably acetyl-CoA). The corresponding 2Fo-Fc electron density map is shown as a blue mesh contoured at 1σ. Superimposed and depicted with thinner sticks, the acetyl-CoA allosteric activator previously characterized on MsSucA$_{D360}$ in the resting state (PDB 2XTA [6]), showing the equivalent pose. To note, in both cases the absence of supporting electron density for the pantothenate chain hindered tracing the ligand in its entirety.

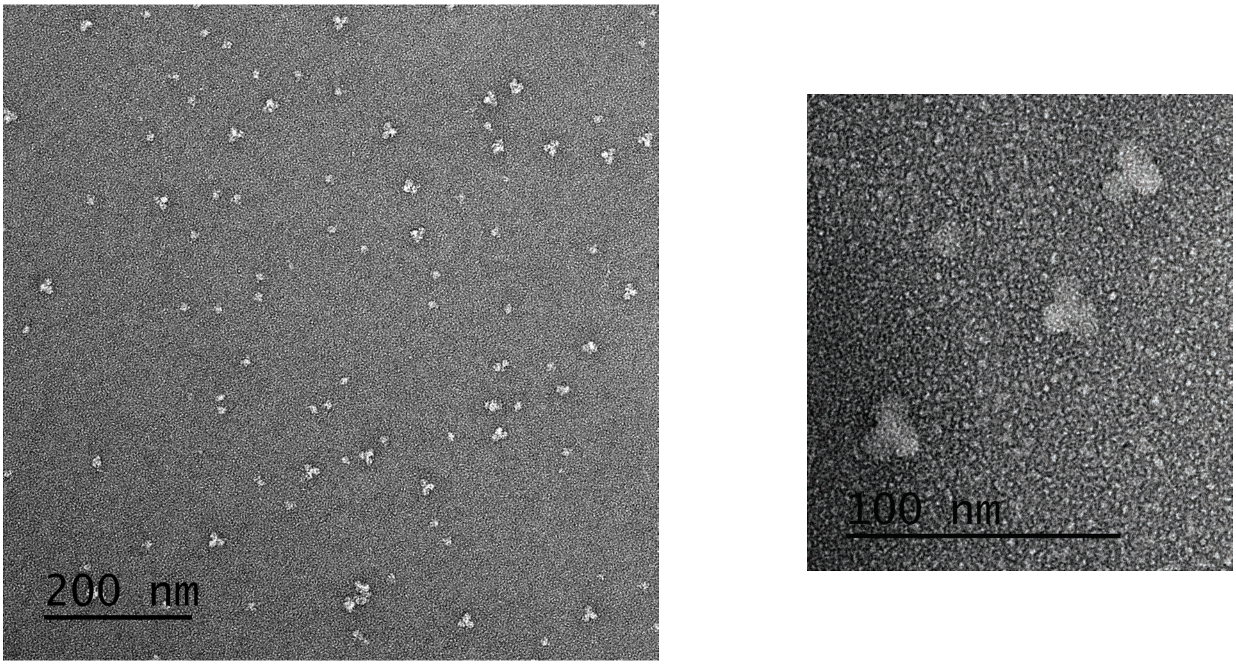


**Suppl. Figure 8. Negative stain EM of an OdhA purified sample.**
Uranyl acetate negative staining of OdhA molecules, from a 0.01 mg/ml solution, observed at magnification of 49000x (left) and 180000x (right). Size and shape of the particles are consistent with the homohexameric $OdhA_{\Delta 97}$ structure determined by X-ray crystallography. Scale bars on the bottom left.

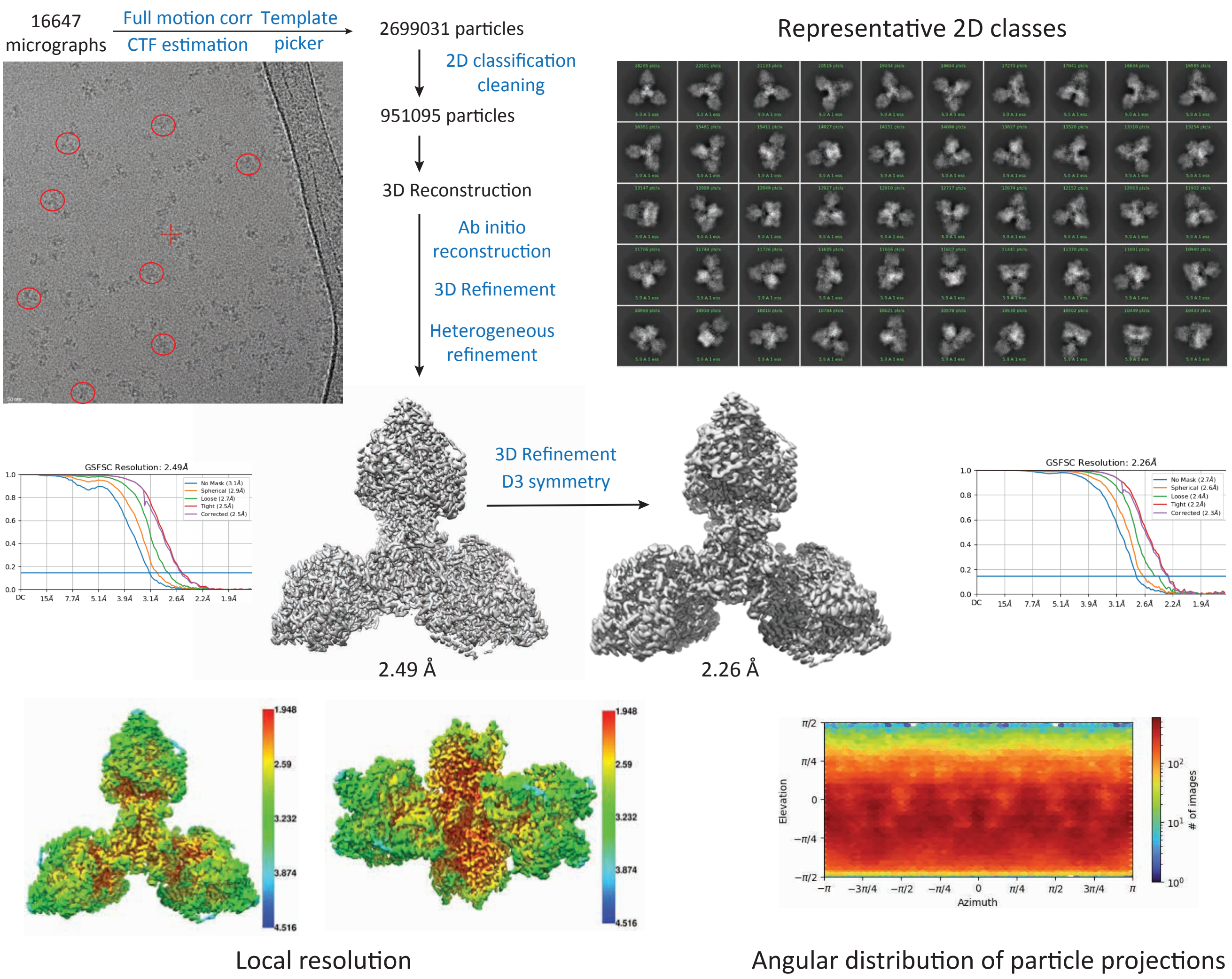


**Suppl. Figure 9. Single particle cryo-EM processing pipeline.**
Scheme representation of the single particle cryo-EM data processing pipeline for the OdhA-SP dataset. Top left: reprensentative micrograph, particles indicated by red ellipses; top right, representative 2D classes; center, EM maps after *ab initio* reconstruction and 3D refinement, before and after further refinement applying D3 symmetry (GSFSC resolution curves at the left and right, respectively). On the bottom, top and lateral view of the final map, coloured according to local resolution (red-blue scale on the right, in Å), and angular distribution of particle projections. Figures retrieved from cryoSPARC [7], map views at the center generated by ChimeraX [8]. All OdhA single particle cryo-EM datasets were processed following the same pipeline (Table 2).

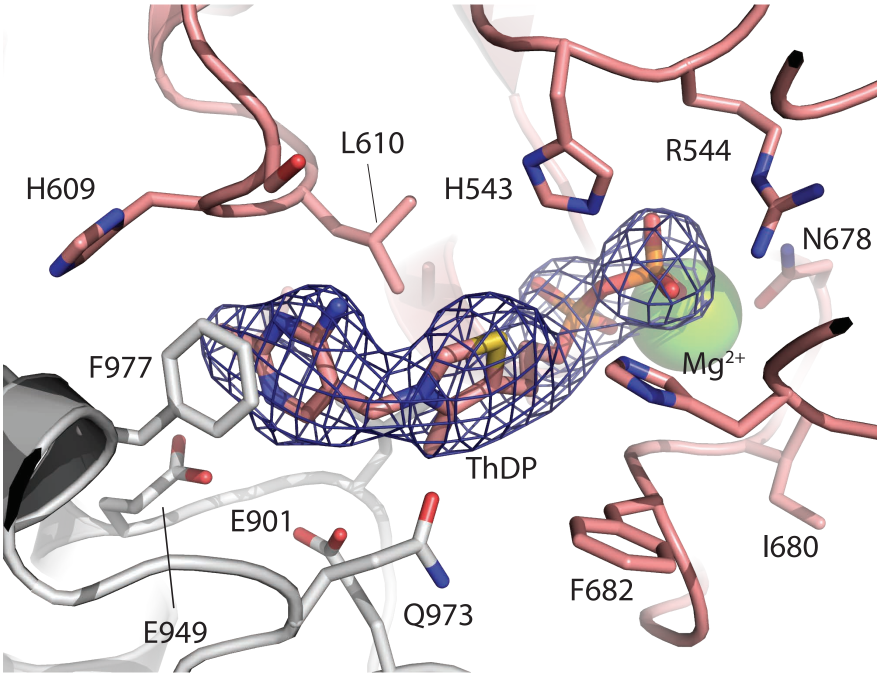


**Suppl. Figure 10. View of the OdhA E1o active site, single particle EM structure.**
Zoomed view on the E1o active site (cartoon representation) of OdhA, determined by single particle EM. The ThDP cofactor is shown in sticks, $Mg^{2+}$ as a green sphere. The EM map for the ligand is shown as a blue mesh and contoured at the 3σ level. The two OdhA protomers contributing to the active site are depicted in pink and gray, respectively. Side chains of residues in close proximity to $Mg^{2+}$-ThDP are shown.

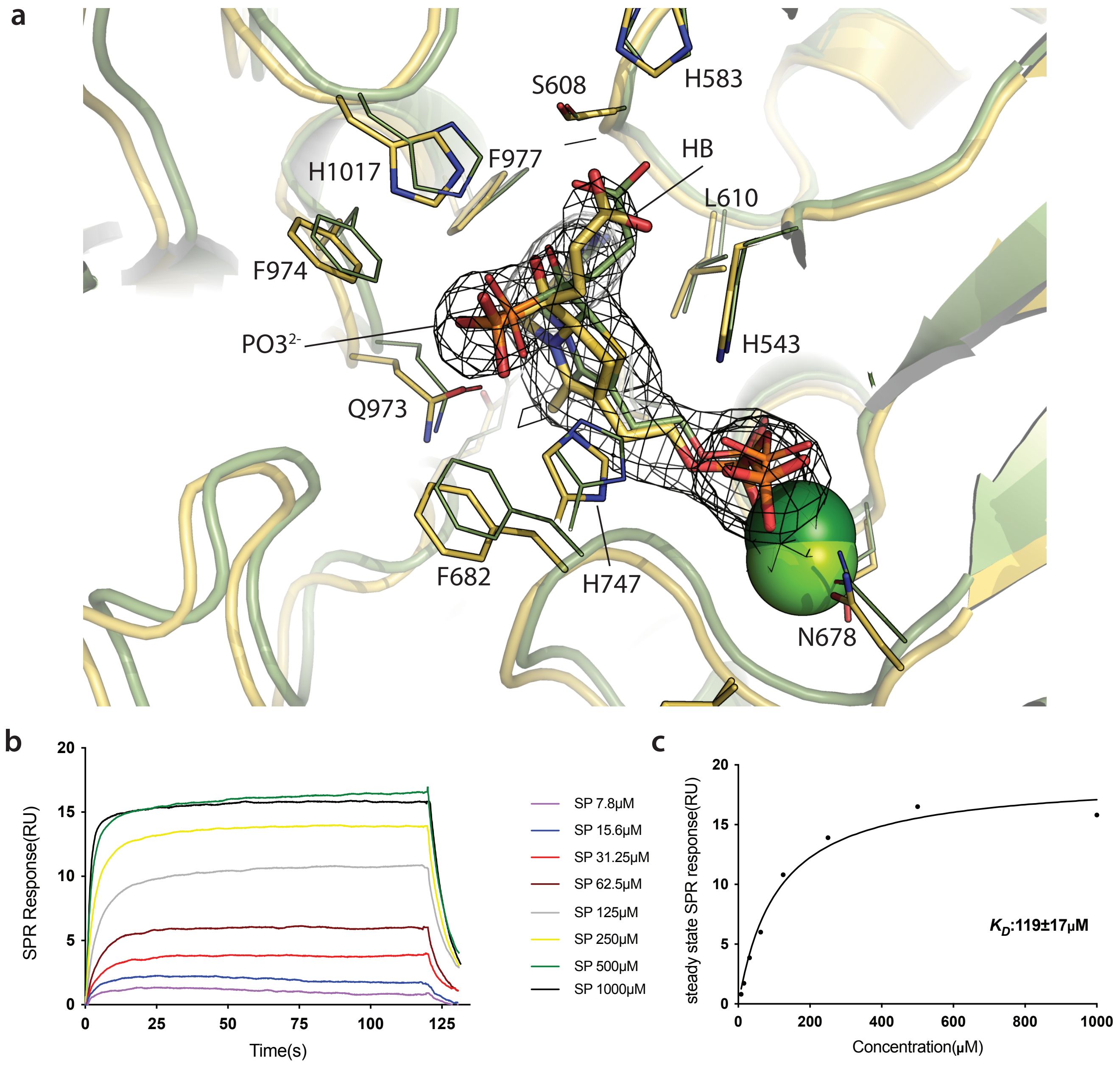


**Suppl. Figure 11. OdhA E1o active site following reaction with succinyl phoshonate (SP).**
(**a**) View of the OdhA active site (cartoon, yellow) following reaction with succinyl phosphonate, superimposed to *Ms*KGD$_{\Delta 360}$ cocyristallized with SP (green; PDB 6R29 [9]). The mesh corresponds to the density modified EM map for the ThDP-SP covalent adduct, contoured at the $3\sigma$ level. Note the equivalent orientation of both the phosphonate group, approximately perpendicular to the thizaolium plane, and the hydroxybutyryl moiety (HB) with respect to the corresponding adduct in *Ms*KGD [9], as well as the overall conservation of the active site environment. Active site residues indicated following OdhA numbering. (**b**) Surface plasmon resonance sensorgram of SP (concentrations indicated on the right) flown over $His_6$-tagged OdhA bound to the chip surface. (**c**) Steady state fitting from surface plasmon resonance experiments. The calculated $K_D$ value is reported.

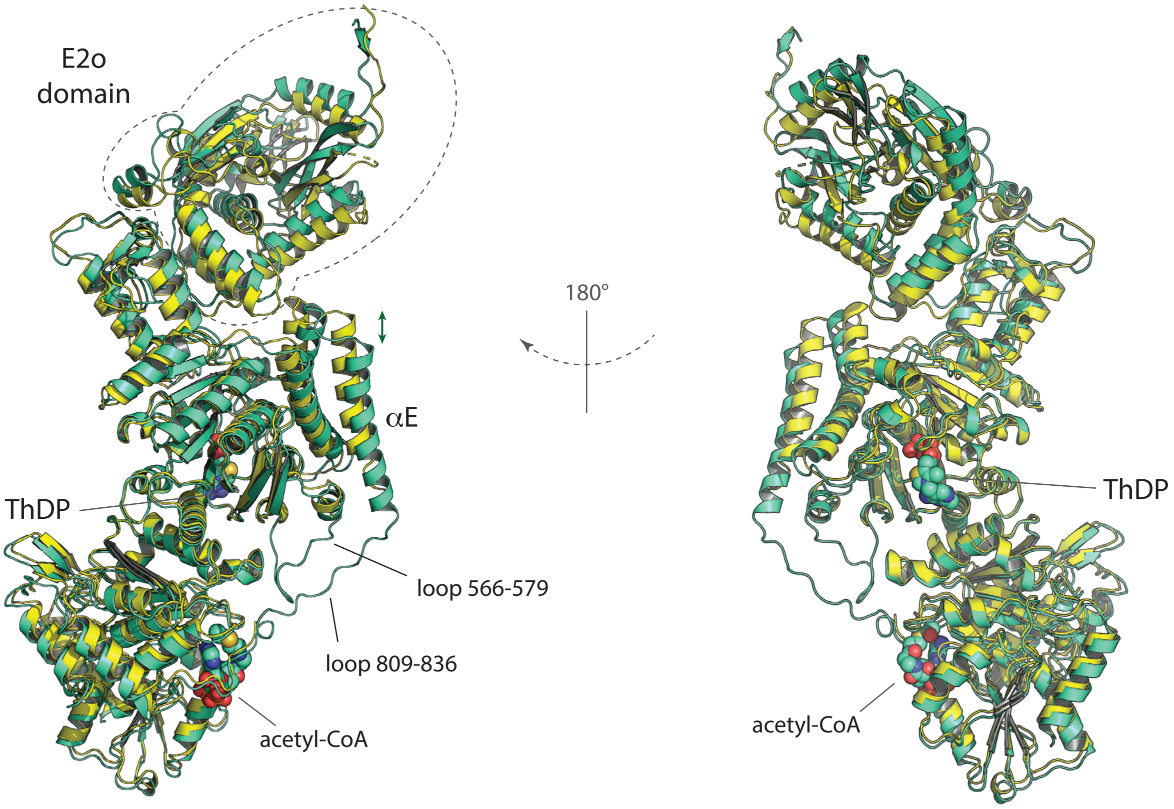


**Suppl. Figure 12. Comparison of OdhA protomers from X-ray crystallography and single particle cryo-EM.** Cartoon view of superimposed OdhA models obtained from X-ray crystallography (OdhA$_{\Delta97}$, 2.5 Å resolution; yellow) and single particle EM (2.2 Å resolution; green), limited to a single OdhA monomer. The E2o domain is located on the top (dashed contour). To note the different enzyme conformations, *i.e.* resting (OdhA$_{\Delta97}$) *vs*. active (EM model). The conformational change of the E1o domain can be simplified via the αE helix deplacement (green arrow) and the disorder to order transition of loops 566-579 and 809-836. The ThDP cofactor and the acetyl-CoA allosteric activator, as bound in the EM structure, are depicted as spheres with van der Waals nonionic radii. The rmsd following the superimposition of the E1o domains is 0.94 Å.

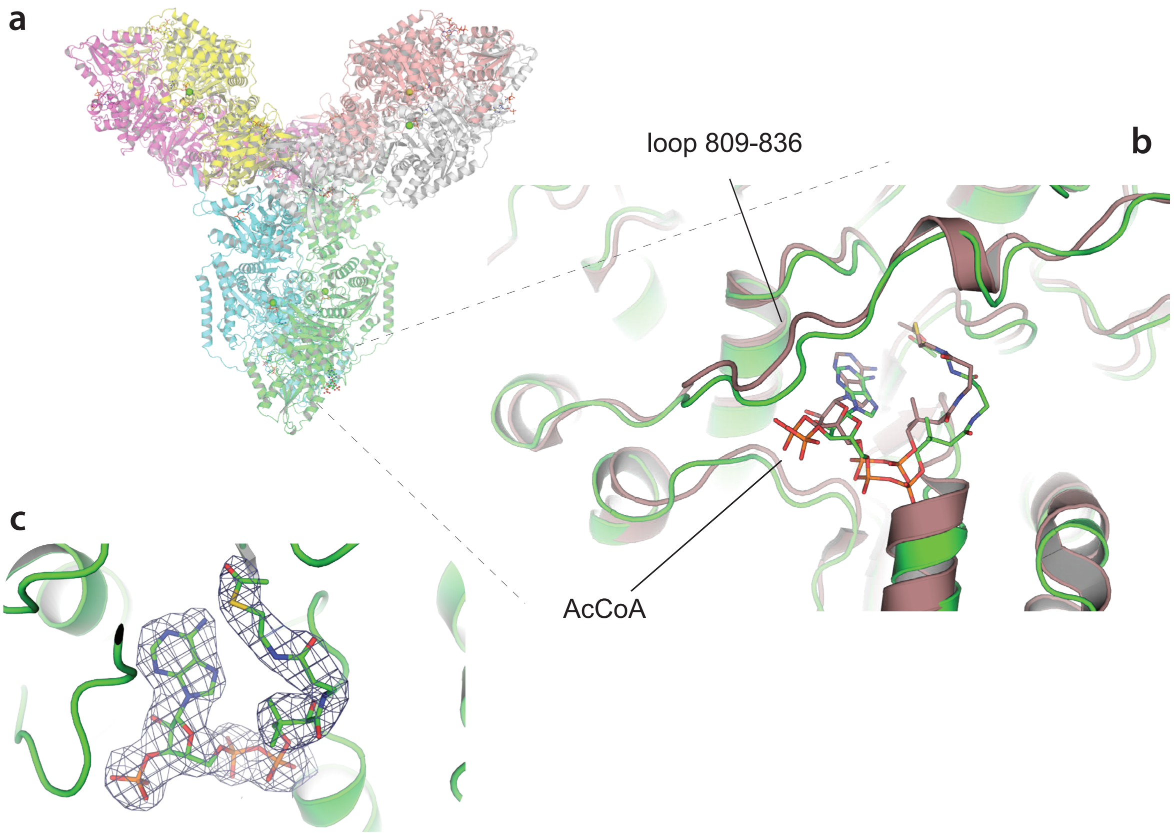


**Suppl. Figure 13. OdhA acetyl-CoA allosteric binding site in single particle cryo-EM models.**
(**a**) Cartoon representation of hexameric OdhA, determined by single particle cryo-EM (one colour per chain) and zoomed view (b) on the allosteric acetyl-CoA binding pocket (green and light blue corresponding to different OdhA protomers), superimposed to the structure of *Ms*SucA$_{\Delta 360}$ in complex with enamine-ThDP and acetyl-CoA (brown; PDB 2Y0P [6]). Note the equivalent pose of acetyl-CoA bound to OdhA (balls and sticks, color per atom) and to *Ms*SucA$_{\Delta 360}$ (brown sticks). Indicated is the loop 809-836 (OdhA numbering; green), connecting the first to the second $\alpha/\beta$ domain within the E1o domain, and already shown in *Ms*SucA to be stabilized by acetyl-CoA binding [6]. (c) EM density for the bound acetyl-CoA in the OdhA model (no added ligands; 2.2 Å resolution), contoured at the 5$\sigma$ level. All single particle EM models of OdhA, with the notable exception of the OdhA-OdhI complex, show acetyl-CoA bound in the same position.

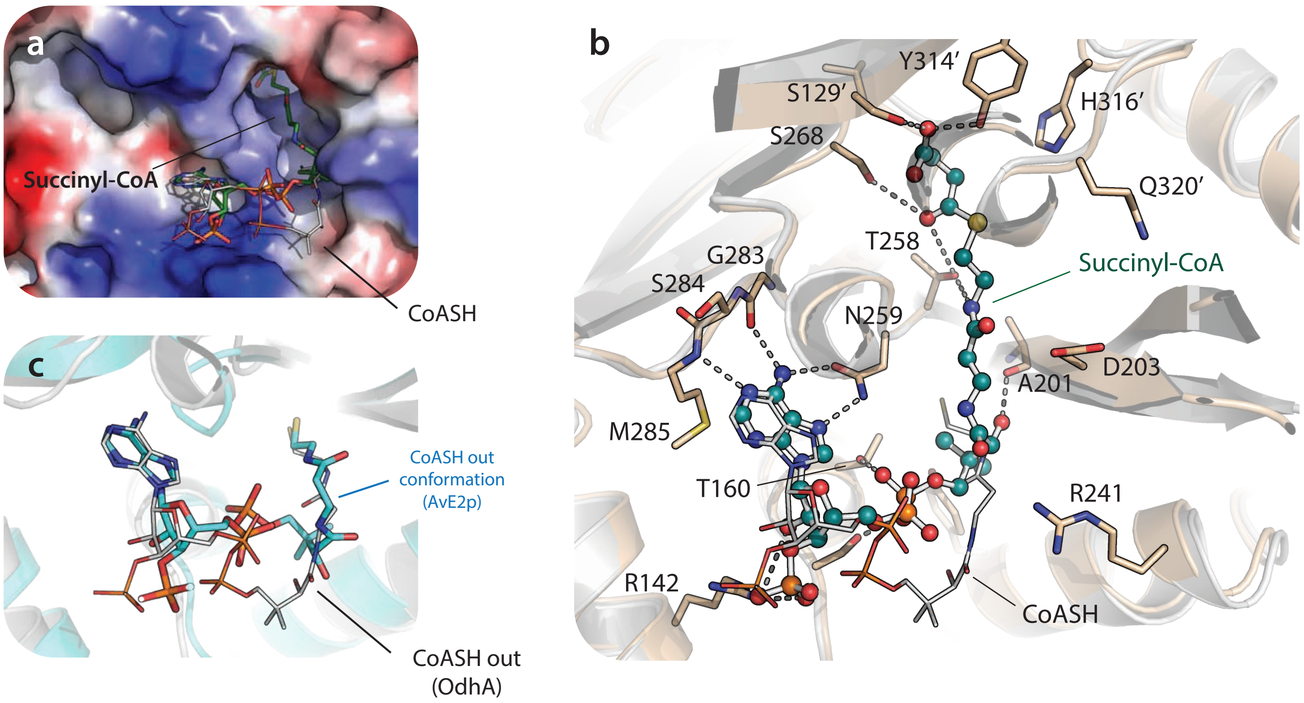


**Suppl. Figure 14. CoA esters binding to the OdhA E2o domain.**
(**a**) Comparison of CoASH and succinyl-CoA binding to the OdhA E2o domain surface, colored by its electrostatic potential (calculated by pymol [5]). To note the distinct conformation of the pantothenate chain in the two cases, with succinyl-CoA entering the tunnel to the active site. (**b**) Cartoon top view of the same superpostion as in (a); the OdhA-CoASH complex is depicted in gray, while the OdhA-succinyl-CoA complex is shown in light brown. Residues involved in ligand binding or with a predicted catalytic role (i.e. H316′, Q320′) are shown. Grey dashed line indicate polar interaction between OdhA residues and succinyl-CoA. (**c**) Comparison of the CoASH pose on the OdhA E2o domain (cryo-EM structure, this work) and the observed CoASH out conformation described for *A. vinelandii* E2p [10].

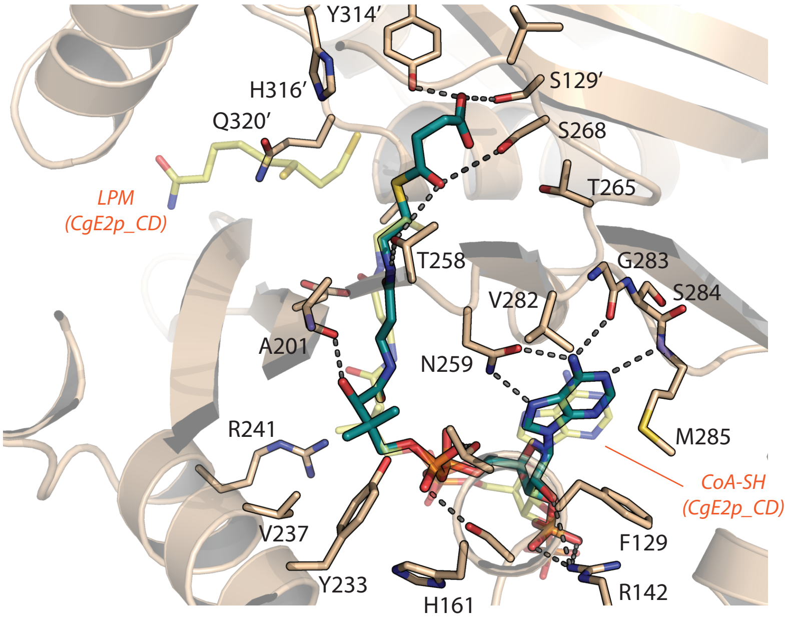


**Suppl. Figure 15. Succinyl-CoA binding to the E2o domain.**
Detailed view of the OdhA E2o active site from the OdhA-succinyl-CoA complex. The bound succinyl-CoA is depicted as sticks (atoms in green); hydrogen bonds between the enzyme the ligand are shown as dashed, gray lines. On the background (yellow, semi-transparent sticks) CoASH and lipoamide extracted from the ternary complex of *C. glutamicum* E2p (PDB 6ZZK; [11]), following superimposition of the entire E2p trimer to the OdhA E2o domain (superimposed E2p protein model not shown). To note the very close pose of CoASH and succinyl-CoA in the two proteins.

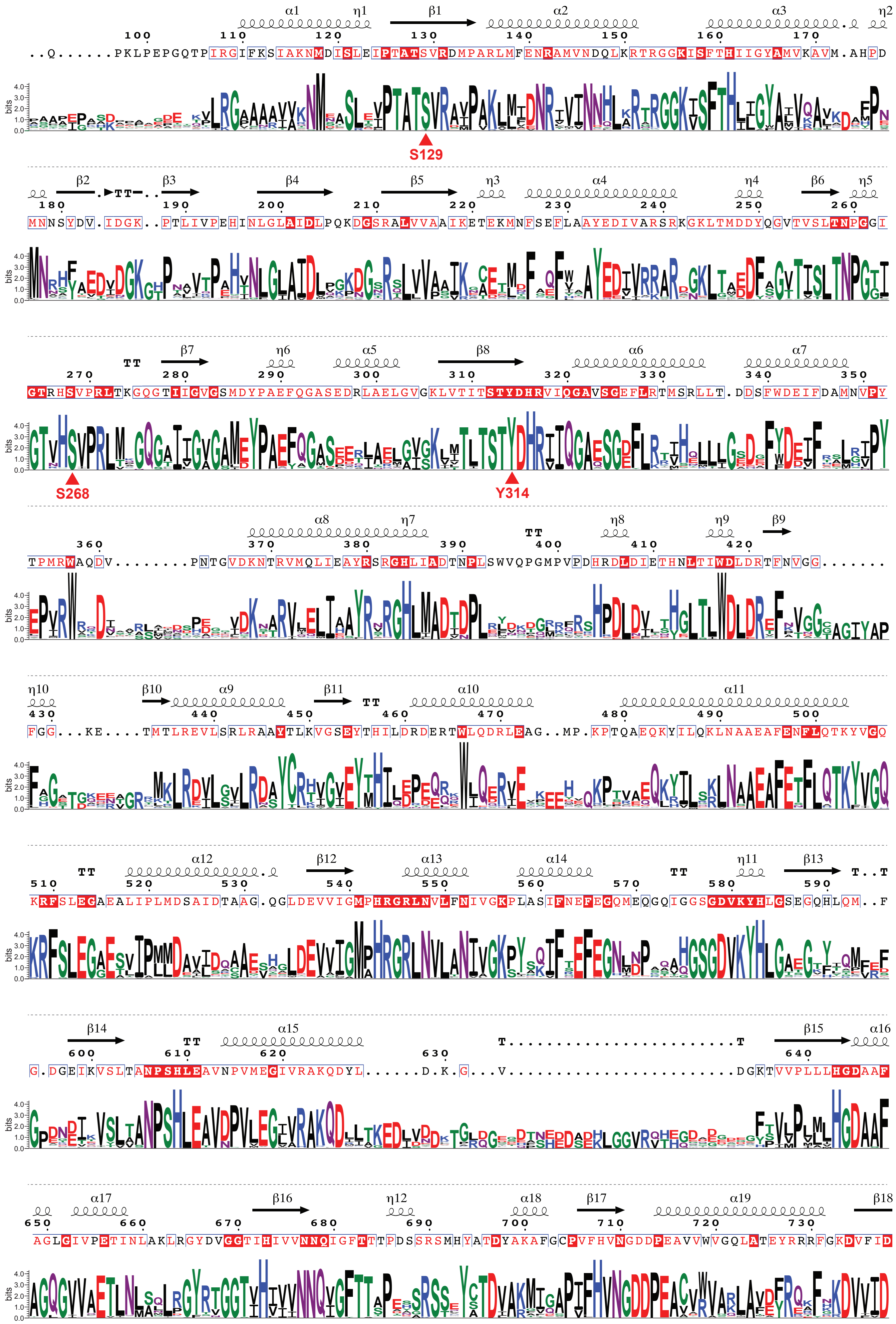
S129
S268
Y314

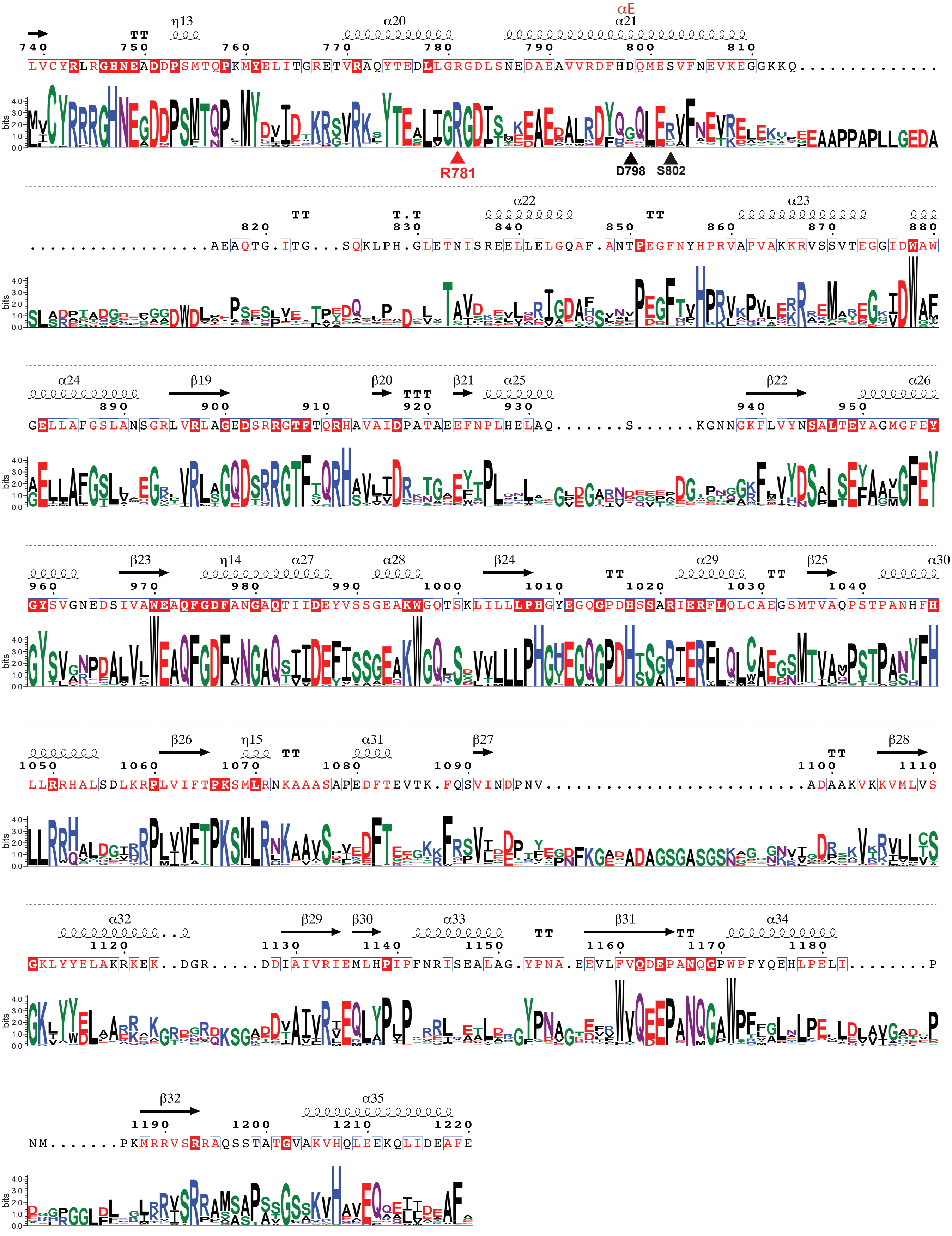

**Suppl. Figure 16. Logo reprensentation of multiple sequence aligment of actinobacterial OdhA orthologues.**
Sequence logo based on a multiple sequence alignment (MSA) of OdhA from representative members of the Actinobacteria phylum; the analysis is here limited to OdhA residues 95-1220, excluding the N-terminal, flexible portion. The overall height of each stack indicates the sequence conservation at that position, and the height of the symbols within each stack indicates the relative frequency of each amino acid at that position. Positively charged amino acids are indicated in green, negatively charged amino acids in blue and amino acids with hydrophobic side chains in red. On top of the logo, the sequence of *C. glutamicum* OdhA is indicated, as well as its secondary structure inferred from the single particle cryo-EM model of OdhA (no added ligands). A position is framed in blue if more than 70% of the residues in the MSA are similar according to physicochemical properties. A position is boxed in red if all residues in the column are identical, and a position has a red font is the residues are similar. α-helices are displayed as squiggles, β-strands are rendered as arrows, strict β-turns as TT letters and strict α-turns as TTT. OdhA primary sequence and secondary structure were depicted with ESPpript [4]. Arrows point to OdhA residues highlighted in the main text (red: conserved, black: not conserved).

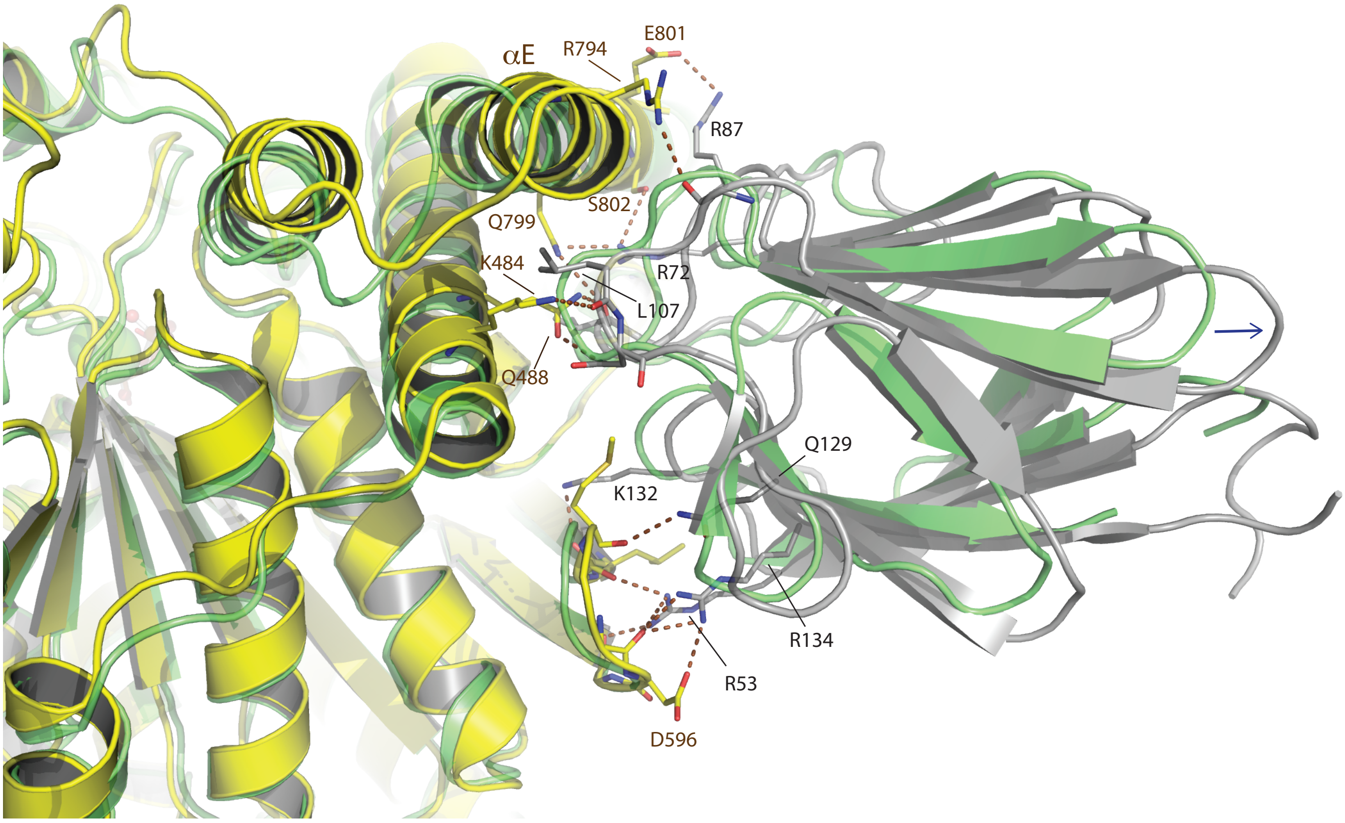


**Suppl. Figure 17. OdhA-OdhI interactions.**
Superimposition of the OdhA-OdhI complex, solved by single particle cryo-EM (OdhA depicted in yellow, OdhI in grey), to the *Ms*KGD$_{\Delta 360}$-GarA complex (PDB 6I2Q [1]; green). Side and/or main chain atoms from residues involved in protein-protein interactions are shown in sticks; brown dashed lines indicate polar interactions (hydrogen bonds, salt bridges). For clarity, only intermolecular OdhA-OdhI interactions are depicted. To note, OdhI (grey) is shifted ~ 2.2 Å (calculated as rmsd on C$\alpha$) aside from the GarA binding position (green; indicated by the arrow), despite the close superimposition of the E1o domain core between OdhA and *Ms*KGD. OdhI and GarA can otherwise be superimposed within an rmsd of 0.5 Å, confirming the overall conservation of the $\beta$-sandwich fold characteristic of FHA domains.

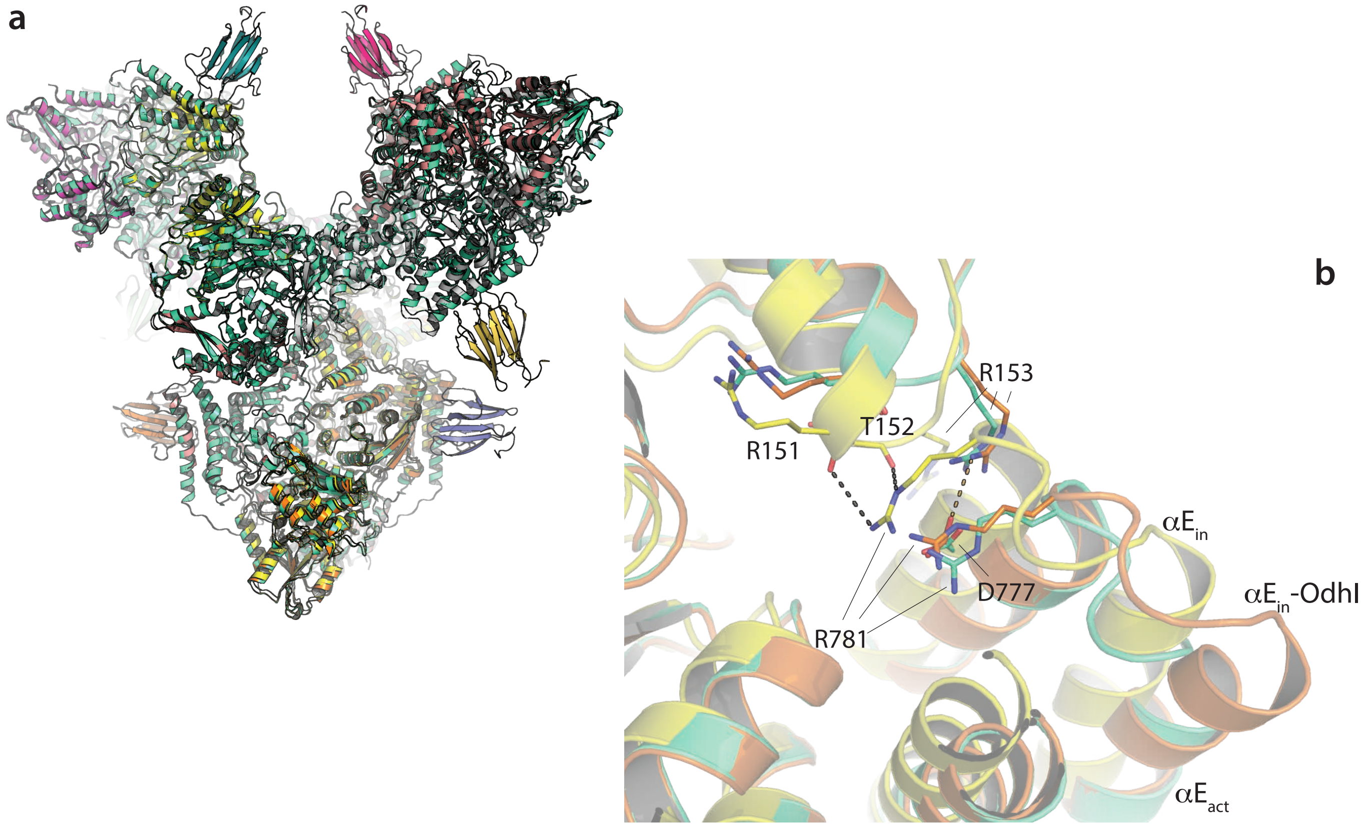


**Suppl. Figure 18. E2o-E1o interdomain interactions in OdhA.**
(**a**) Superimposition of the OdhA$_{\Delta 97}$ (crystal structure; yellow), OdhA-OdhI (single particle cryo-EM structure, resting conformation; coloured by chain) and OdhA alone, also solved by single particle cryo-EM (active conformation; teal colour). (**b**) Zoom on the interface region between the E2o and E1o domains, at the top of helix $\alpha$E. To note the different position of the loop preceeding $\alpha$E and bringing the conserved Arg781, which H-bonds to Arg151 and Thr152 main chain carbonyl oxygens only in the OdhA$_{\Delta 97}$ model (similarly to previous observations on *Ms*KGD [6]). In all the cryo-EM OdhA structures described here, regardless of the E1o domain conformation (resting *vs*. active), a salt bridge between Asp777 and Arg153 is observed instead.
